# Layered Feedback Control Overcomes Performance Trade-off in Synthetic Biomolecular Networks

**DOI:** 10.1101/2021.09.12.459953

**Authors:** Chelsea Y. Hu, Richard M. Murray

## Abstract

Layered feedback is an optimization strategy in feedback control designs widely used in electrical and mechanical engineering. Layered control theory suggests that the performance of controllers is bound by the universal robustness-efficiency tradeoff limit, which could be overcome by layering two or more feedbacks together. In natural biological networks, genes are often regulated with redundancy and layering to adapt to environmental perturbations. Control theory hypothesizes that this layering architecture is also adopted by nature to overcome this performance trade-off. In this work, we validated this property of layered control with a synthetic network in living *E. coli* cells. We performed system analysis on a node-based design to confirm the tradeoff properties before proceeding to simulations with an effective mechanistic model, which guided us to the best performing design to engineer in cells. Finally, we interrogated its system dynamics experimentally with eight sets of perturbations on chemical signals, nutrient abundance, and growth temperature. For all cases, we consistently observed that the layered control overcomes the robustness-efficiency trade-off limit. This work experimentally confirmed that layered control could be adopted in synthetic biomolecular networks as a performance optimization strategy. It also provided insights in understanding genetic feedback control architectures in nature.

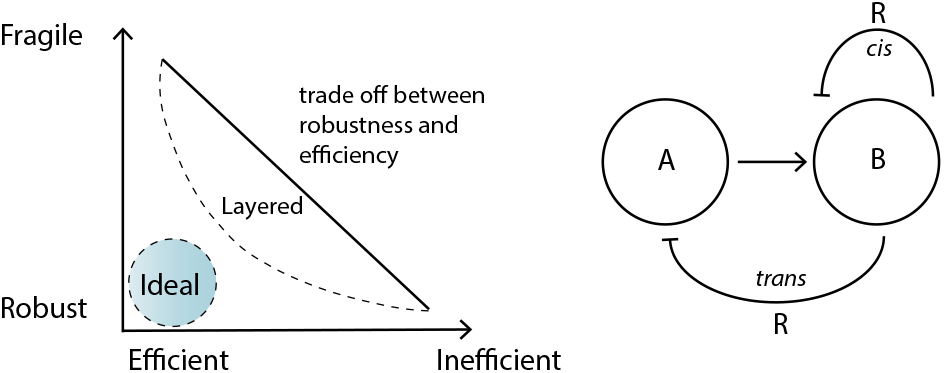

## Introduction

Two of the major goals of synthetic biology are to engineer biological systems to perform desired tasks and to understand biomolecular networks by building them. These two goals are both very ambitious; fortunately, their advancements are strongly coupled. By building and studying synthetic systems, we gain insights into the natural networks; with a deeper understanding of how natural genetic networks allow life to persevere through constant adversity, we develop stronger theoretical principles to guide synthetic network design. Like many other systems humans have engineered, feedback controls are essential components of a reliable synthetic biological system. Over the past two decades, the synthetic biology toolbox of characterized genetic components, modules, and motifs has been expanding exponentially [1]. As the complexity of synthetic circuits grows, its interplay with control theory becomes more prominent. A significant amount of work has already shown that applying feedback controls improves the reliability, efficiency, and performances of synthetic biological systems [2, 3, 4, 5, 6, 7, 8]. Feedback control provides correcting actions based on the difference between desired and actual performance [9, 10], therefore it buffers systems from external disturbances and variations of components within the system. However, feedback can also destabilize the system when improperly designed [10]. Feedback design is especially challenging in biological systems due to its complexity — all molecular species are part of an extensive endogenous network that consists of numerous feedback mechanisms.

In nature, biology has developed remarkably sophisticated strategies to apply feedback controls to biomolecular and physiological networks. Interestingly, layering and redundancy appears to be a common style in these feedback architectures. For instance, bacteria cells layer a positive control to move towards nutrients and a negative control to move away from toxins through chemotaxis signaling [11, 12]. During a heat shock, the endogenous control in bacteria maintains the amount of heat shock proteins with a multi-layer control strategy via translation, chaperone interaction, and protein degradation [13]. At the physiological level, the human body maintains a relatively constant glucose level in the bloodstream through insulin, the production of which is regulated through the interplay of the pancreas with the brain, liver, gut, as well as adipose and muscle tissues [14]. It has also been discovered that the sleep and arousal states of animals are controlled with a layered architecture [15]. Control theorists found that layering is a powerful optimization method for feedback control design [16]. This is because the performance of feedback controls is often bound by hard limits. A system that is optimized for one type of disturbance is typically fragile to other types of disturbance [10, 17]. These constraints are also manifested as the robustness-efficiency trade-off limit, which are theorized to be fundamental and universal for all feedback systems [18, 19]. If a system responses to changes in inputs quickly, it is considered *efficient*; if a system shows strong attenuation against disturbances, it is considered *robust*. This trade-off limit prevents a system from being both *efficient and robust*. It is also theorized that, in both natural and engineered systems, if multiple control modules of various robustness-efficiency profiles are layered together, this hard limits could be overcome [20, 21] (Figure 1A).

**Figure 1:**
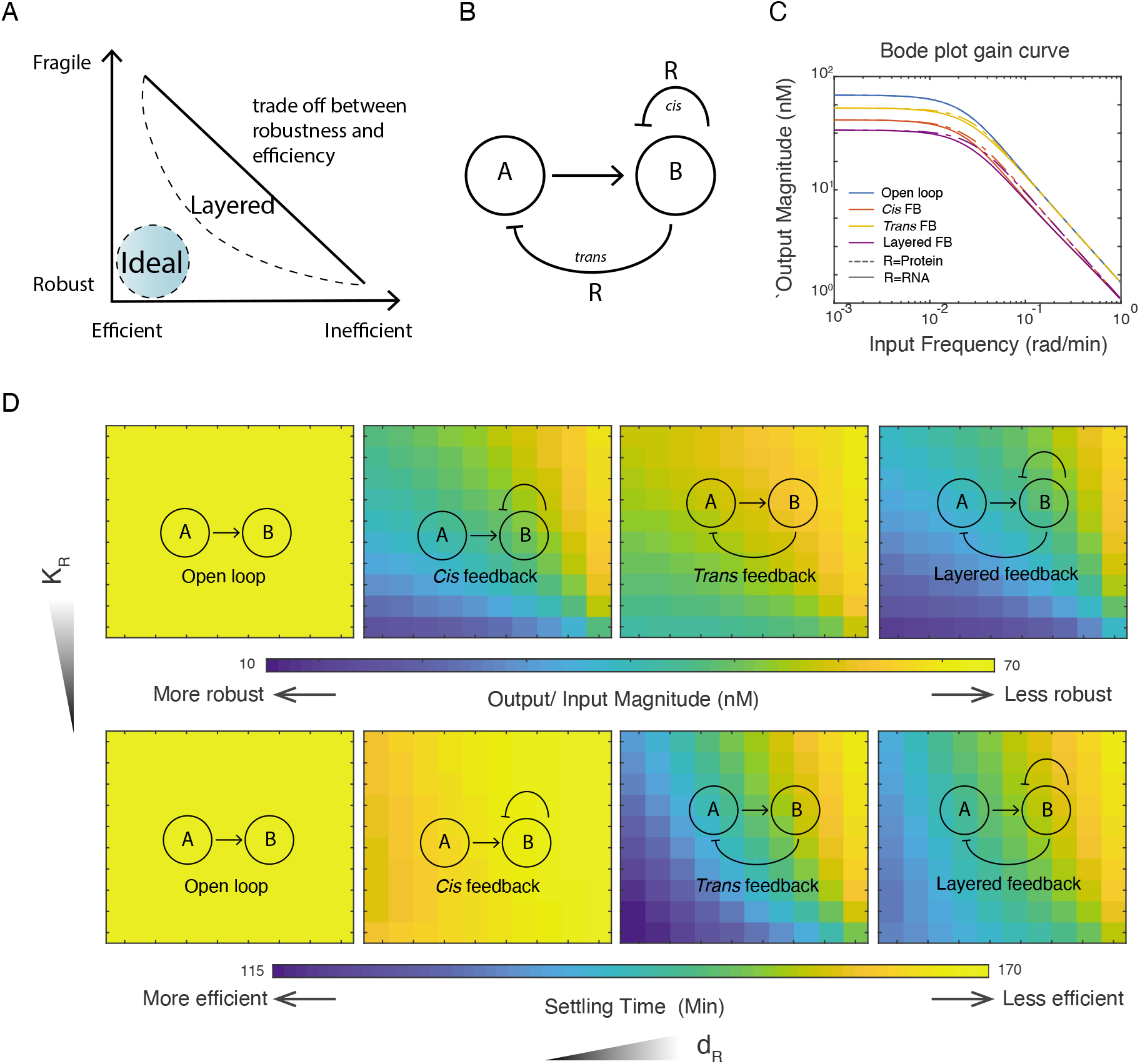
Stability analysis of the layered feedback controller using a minimal model. (A) The theorized law of robustnessefficiency trade-off limit of feedback control. Layering is an optimization strategy to overcome these trade-off limits. (B) Schematics of the node-based system. Here A and B are two molecular species, where A activates the expression of B and R. Functioning as a regulator, R regulates species B directly through the *cis* feedback on B and indirectly through the *trans* feedback on A. When both types of feedback exist in the same system, the configuration is termed layered feedback. (C) Response magnitude with production disturbance at different frequencies. This plot shows the magnitude response of species B (y-axis) when all three species’ production rates are subjected to a perturbation (on *γ*) at different frequencies (x-axis). The blue, red, yellow, and purple curves represent the open loop, the *cis* feedback, the *trans* feedback, and the layered feedback circuit structures, respectively. Solid lines represent the magnitude response of these four structures when regulator R is an RNA species, dashed lines when R is a protein species. (D) Disturbance attenuation and settling time of four architectures in a two-dimensional parameter space *K_R_* and *d_R_*. Top panel shows the output disturbance magnitude when all three species’ production rates are subjected to a perturbation at low frequency. Bottom four panels show the time it takes for the systems to settle back to equilibrium after the perturbation. Both response magnitude and settling time are represented by the color schemes (see color bar). For each of the eight heat maps, the x-axis (*d_R_*) represents the degradation rate of regulator R; a large *d_R_* models a regulator with fast degradation rate. The y-axis (*K_R_*) represents the repression constant of regulator R; a large *K_R_* models a regulator with weak repression strength.

In this work, we forward engineer layered feedback control in living *E. coli* to study the dynamical performance of this architecture in a biomolecular context. We first start with a node-based design of two layers of feedback (Figure 1B) and a minimal model of three species (Equations 1–3). By performing system analysis on the minimal model, we observe the robustness-efficiency trade-off on the two single-layer designs, where one appears to be robust but inefficient while the other is efficient but fragile. Meanwhile, we see a clear integration of these two traits in the layered feedback system. Subsequently, we expand this design into a generic biomolecular model that describes the coarse-grained dynamics of a biomolecular feedback system. With simulation, we also observe that for all four sets of the proposed designs, the two single-layer feedback are bound by the robustness-efficiency trade-off limit, while the layered feedback design overcomes it. Based on the simulated result, we choose the best performing layered feedback design and construct it in *E. coli*. Finally, we present eight sets of dynamical perturbation experiments, with four different types of disturbances, each on two directions. We show that the layered feedback design in all cases overcomes the robustness-efficiency trade-off limit. This result applies layered feedback control in a biomolecular network and validates a key hypothesis of layered control theory. In addition to providing a validated optimization strategy to genetic network engineering, we hope the insights it provides will improve our understanding of natural dynamical systems in biology.

## Results and Discussion

### Robustness-efficiency Trade-off Analysis with a Node-based Design

First, to simplify the problem, we proposed a layered negative feedback control architecture with a node-based design for system analysis. As shown in Figure 1B, we defined two nodes, A and B, where B is the observable output of the system that is activated by A. Species R is a byproduct of species B that negatively regulates the expression of B and itself through two possible routes: *cis* feedback (R represses B) and *trans* feedback (R represses A). The activation and repression here are estimated with first-order Hill functions:

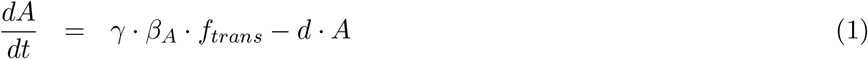

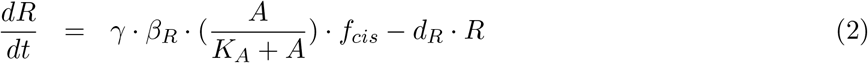

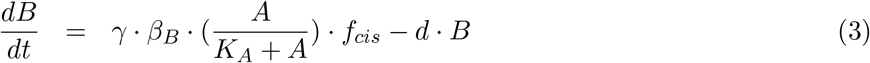

For the open loop, *f_cis_* = *f_trans_* = 1; for the *cis* only feedback, 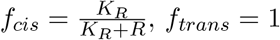; for the *trans* only feedback, 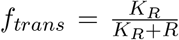; for the layered feedback, 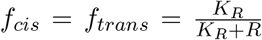. Both A and B are assumed to be protein species, while R could be either protein or regulatory RNA. In the ODEs, *β_A_, β_R_*, and *β_B_* denote the expression rate of A, R, and B, respectively. The constant *K_A_* defines the activation coefficient of A, and *K_R_* defines the repression coefficient of R. Both species A and B degrade at the same rate *d*, while R degrades at rate *d_R_*. We define that if R is a protein species, then *d_R_* = *d*, if R is an RNA species, then *d_R_* = 10 · *d* for faster degradation. To ensure that the protein and the RNA regulator have compatible regulating effectiveness, we constraint the product of *K_R_* · *d_R_* to be constant (i.e. for an RNA regulator *K_R_* = 10, *d_R_* = 0.3, and for a protein regulator *K_R_* = 100, *d_R_* = 0.03). Here we define the output as species B. The input is defined as *γ*, a unitless scalar that impacts the expression rate of all three species (*β_A_* and *β_B_*). Our analysis in the rest of this work focuses on four designs: open loop, the *cis* feedback, the *trans* feedback, and the layered feedback.

In figure 1C, we obtained the system’s Bode plot gain curve to describe the design’s system dynamics. Bode plot is a common tool used in control theory to visualize a linear system’s dynamics in respect to the input’s frequency [10]. Since all four systems are nonlinear, we linearized them at their equilibrium points before converting them to the frequency domain. In this plot, the x-axis represents the input frequency in rad/min, and the y-axis represents the output/input magnitude, which has a unit of nM. When the input frequency is low, this could be understood as a long-lasting perturbation to the universal species production scalar γ. When the input frequency is high, the input signal oscillates rapidly, and the output response to the disturbances diminishes. In this figure, we observe that all three types of feedback attenuate disturbances at low input frequency. In which, the *cis* feedback (red) has a stronger attenuation effect than the *trans* feedback (yellow), and the layered feedback shows the strongest attenuation effect (purple). As the input frequency increases, the *trans* feedback loses disturbance attenuation property compared to the open loop. However, the *cis* feedback control remains more robust than the open loop in the entire frequency span. We also observed that there is only a slight advantage for protein regulator (dashed lines) relative to an RNA regulator (solid lines) at intermediate input frequencies.

To investigate the impact of species R on the system dynamics, we analyze the disturbance magnitude and settling time of the system in a 10 × 10 parameter space that defines the property of regulator R. A large *K_R_* (y-axis) models a weak regulator, and a large *d_R_* (x-axis)models a fast degrading regulator. RNA species usually degrade faster than protein species. We found that over a wide parameter space, the *cis* feedback attenuated disturbances at low frequency better than the *trans* feedback, while the layered feedback showed the most attenuation. In each feedback design, regulators with strong repressive strength and a slow degradation rate achieved the most disturbance attenuation. In the lower panel of Figure 1C, we plotted the settling time of each construct after a step-disturbance in the same two-dimensional parameter space *K_R_* (y-axis) and *d_R_* (x-axis). We found that although the *cis* feedback had a strong attenuation of disturbance, it took longer to recover back to equilibrium, while the *trans* feedback recovered back to equilibrium faster but attenuated the disturbances less effectively. The layered feedback, on the other hand, appeared to have a moderate settling time and strong disturbance attenuation.

### Robustness-efficiency Trade-off Analysis with Generic Biomolecular Configurations

To actuate the node-based design in the biomolecular context, we proposed four possible designs (Figure 2A) with the regulator species being either R (regulatory small RNA, or sRNA) and P (regulatory protein). As shown in Figure 2A, the system is induced with small molecule *x*, which activates the transcription of protein *P_ind_*. Protein *P_ind_* sequentially activates the gene of interest (GOI) cassette, which contains gene GOI, R, and/or P. The four illustrations in Figure 2A represent four designs of layered feedback: (1) sRNA mediated *cis* feedback layered with protein mediated *trans* feedback, (2) sRNA mediated *cis* feedback layered with sRNA mediated *trans* feedback, (3) protein mediated *cis* feedback layered with sRNA mediated *trans* feedback, and (4) protein mediated *cis* feedback layered with protein mediated *trans* feedback.

**Figure 2:**
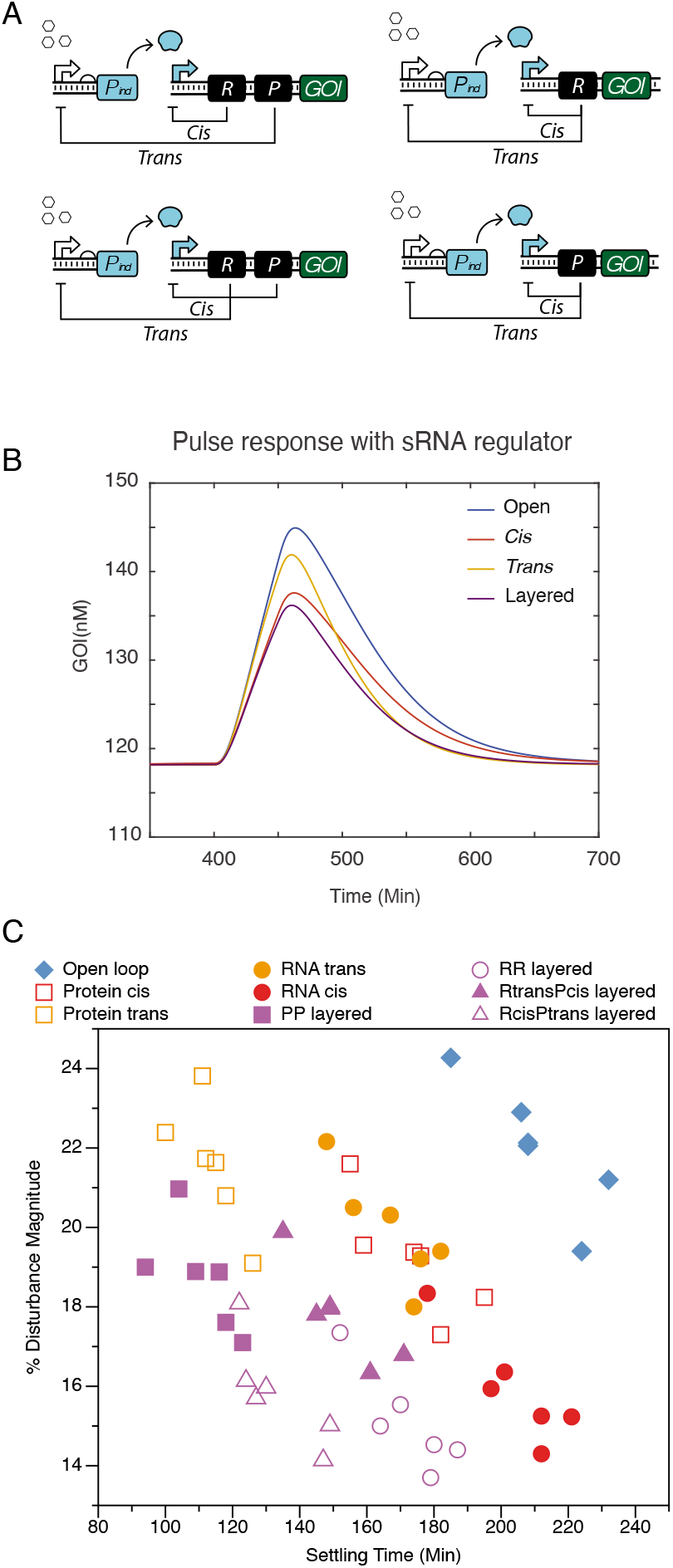
Efficiency-stability trade-off analysis on generic biomolecular configurations. (A) The four possible biomolecular configurations of the node-based design as shown in Figure 1B, with the regulator species being either sRNA or protein. In all cases, gene regulations happen at the transcriptional level. (B) The simulated dynamics of system with a transcriptional pulse perturbation at steady state. This plot corresponds to one of the four designs, where sRNA regulates via both *cis* and *trans* feedbacks. (C) The robustness-efficiency trade-off analysis of nine configurations. The plot is generated from simulated dynamics with randomized model parameters, one example of which is shown in 2B.

Next, we interrogated the performance of these four designs in simulation with pulse disturbances. The dynamics of these systems are described with reduced differential equations (Equation 4–11). The detailed species and parameters involved in this model are listed in Table 1 and Table 2, respectively. It is worth noting that we purposefully chose the degradation rates (*d_r_, d*) and repression coefficients for regulatory RNA *R* and regulatory protein *P*(*K_R_, K_p_*) to simulate the same effective repressive strength for fair comparison. The regulatory RNA is simulated to have strong binding affinity (small *K_R_*) but is quickly degraded (large *d_r_*); the regulatory protein is simulated to have weak binding affinity (large *K_p_*) but is slowly degraded (small *d*).

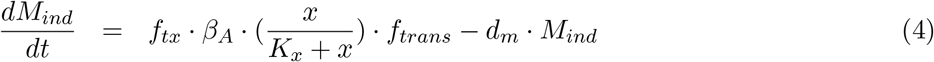

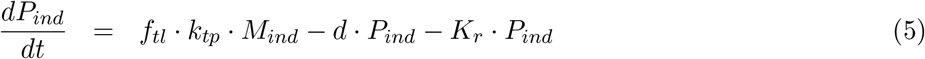

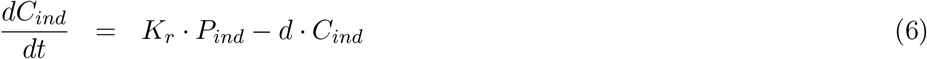

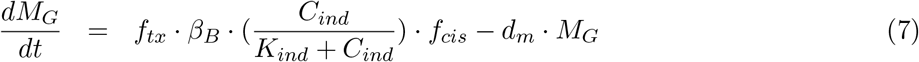

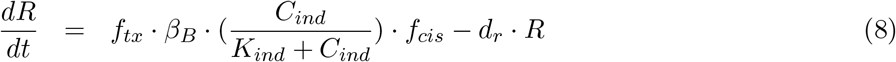

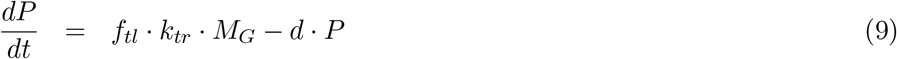

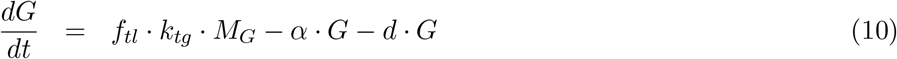

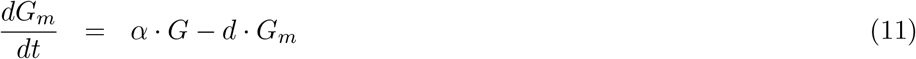

**Table 1:** Generic Biomolecular Model Species

| Species | Description |
| --- | --- |
| $M_{ind}$ | the mRNA of signaling protein $P_{ind}$ |
| $P_{ind}$ | the signaling protein translated peptides |
| $C_{ind}$ | the signaling complex, with folded signaling protein bound with inducer molecules |
| $R$ | the regulator sRNA repressor |
| $P$ | the regulator protein repressor |
| $M_G$ | the GOI mRNA transcript |
| $G$ | the translated GOI peptides |
| $G_m$ | the mature GOI (observable) |

**Table 2:**
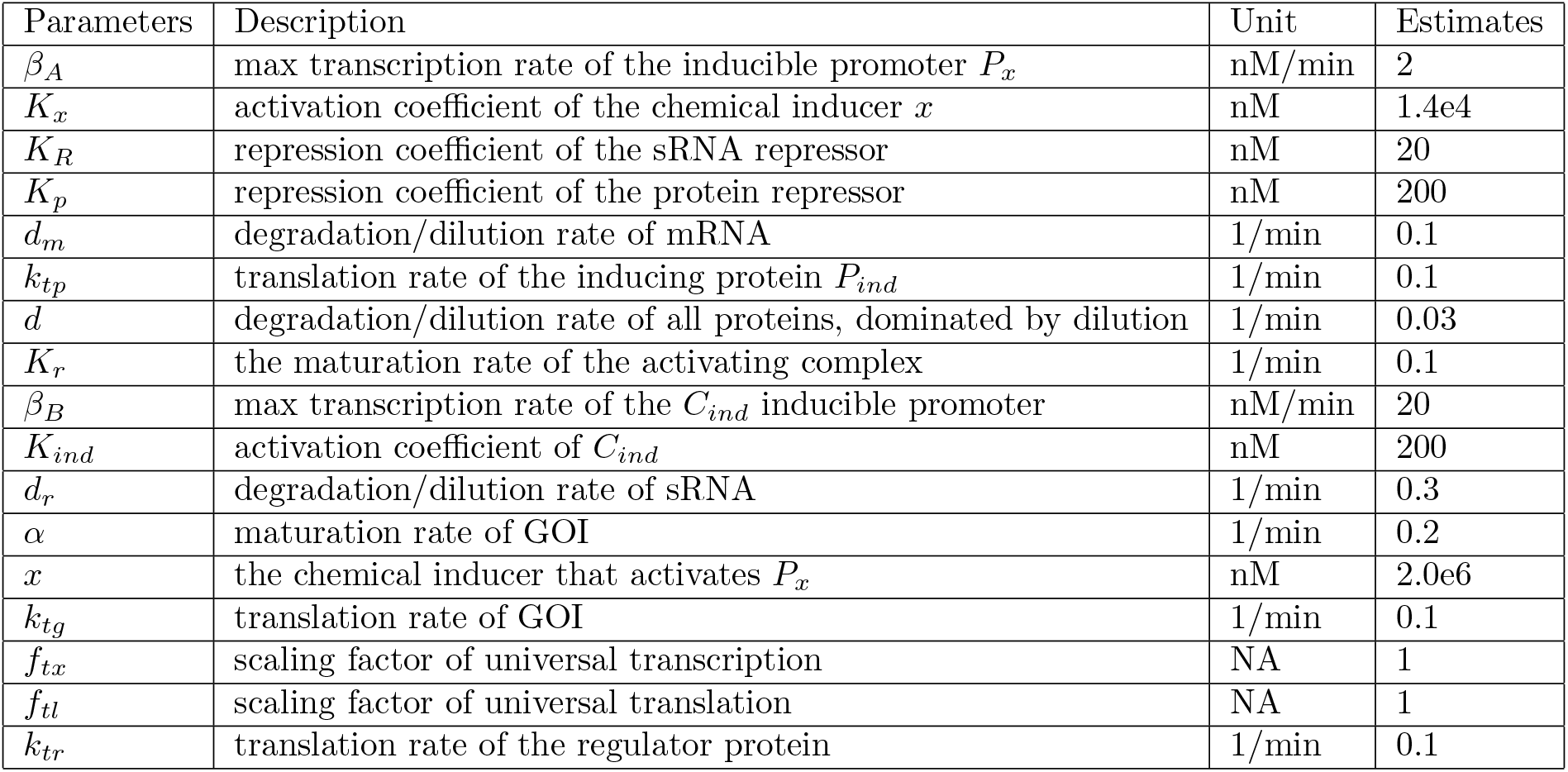
Generic Biomolecular Model Parameters

If transcription is regulated by a protein species, the Hill function is written as 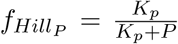. If transcription is regulated by a sRNA species, the Hill function is written as 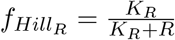. For the open loop, *f_cis_* = *f_trans_* = 1. For the *trans* only feedback, *f_cis_* = 1 and *f_trans_* = *f_Hill_P__*, if mediated by a a protein species; *f_trans_* = *f_Hill_R__*, if mediated by a sRNA species. For the *cis* feedback, *f_trans_* = 1 and *f_cis_* = *f_Hill_P__*, if mediated by a protein species; *f_cis_* = *f_Hill_R__*, if mediated by a sRNA species. For the layered feedbacks, the four possible combinations are: (1) R *cis* R *trans, f_cis_* = *f_Hill_R__*, *f_trans_* = *f_Hill_R__*; (2) R *cis* P *trans*, *f_cis_* = *f_Hill_R__*, *f_trans_* = *f_Hill_P__*; (3) P *cis* R *trans*, *f_cis_* = *f_Hill_P__*, *f_trans_* = *f_Hill_R__*; (4) P *cis* P *trans*, *f_cis_* = *f_Hill_P__*, *f_trans_* = *f_Hill_P__*.

We simulated the system dynamics of all four sets of designs and compared them with their corresponding single-layer and open loop configurations. One of the four sets of dynamics is plotted in Figure 2B. We first scaled the expression of all four configurations to the same equilibrium signal value. At equilibrium, we gave the universal transcriptional rate scalar *f_tx_* a 25% step pulse and held it for 50 minutes. Then we plotted each construct’s percentage GOI response at its peak and the settling time after the disturbance. For a total of nine different configurations (one open loop, two *cis* feedbacks, two *trans* feed-backs, and four layered feedbacks), we generated six sets of parameters with a 25% randomization around the initial guesses for each simulation. Then for each of the nine configurations, the six sets of parameters were used to simulate six dynamical profiles in response to perturbations. The settling times and percentage disturbance magnitude of each simulation were recorded in the robustness-efficiency trade-off plot in Figure 2C. Similar to the result we obtained from the node-based design in Figure 1, we found that the *cis* feedback is more robust against disturbances but recovers slower, *trans* feedback is more fragile against disturbances but recovers faster. As these two feedback controls are layered, the system inherits the strong disturbance attenuation from the *cis* feedback and the fast recovery speed from the *trans* feedback. Finally, we chose the design with sRNA-mediated *cis* feedback and protein-mediated *trans* feedback to proceed to the experimental portion of the work, as it demonstrated the most advantageous performance on the robustness-efficiency trade-off plot.

### Experimental Implementation of a Layered Feedback Control in Living *E. coli* Cells

We designed four genetic constructs to test and experimentally analyze the layered feedback design we proposed based on the simulated results in Figure 2C. We aimed to compare the dynamical performances of the open loop, the *cis* feedback, the *trans* feedback, and the layered feedback. With the effect of genetic context in mind [22], we designed the four constructs with the same configuration. As shown in Figure 3A, the system is composed of two cassettes expressed on two plasmids. The combinatorial promoter *P_Rhl/LacO_* [23] drives the expression of CinR on a medium copy p15A backbone; promoter *P_cin_* drives a another cassette containing sRNA repressor pair (AS and Att) [24], sfYFP, and LacI places on a high copy ColE1 backbone. The system is activated by the induction of combinatorial promoter *P_Rhl/LacO_* with an AHL inducer Rhl. The promoter then allows the expression of protein CinR, which subsequently binds to an AHL inducer Cin in growth medium to activate promoter *P_cin_*. Promoter *P_cin_* controls the transcript that contains sRNA regulating pair AS and Att, the observable system output sfYFP, and the protein regulator LacI. In this configuration, sRNA repressor AS regulates the expression of sfYFP via the *cis* feedback, and LacI represses *P_Rhl/LacO_* to regulate the expression of CinR through the *trans* feedback. To avoid genetic context change and metabolic load variation, we created mutated regulator pieces to disable feedbacks without changing the genetic context. Specifically, we paired AS with its orthogonal attenuator Att(M) [25] to disable the *cis* feedback, and we built LacI(M) off LacI with the LacO binding site sequence removed to disable the *trans* feedback. The construction of these four genetic networks is a non-trivial task. It involves one combinatorial regulation, one activation cascade, and two nested autoregulatory motifs. Extensive part-characterization and expression optimization is required to confirm the networks’ proper dynamical properties.

**Figure 3:**
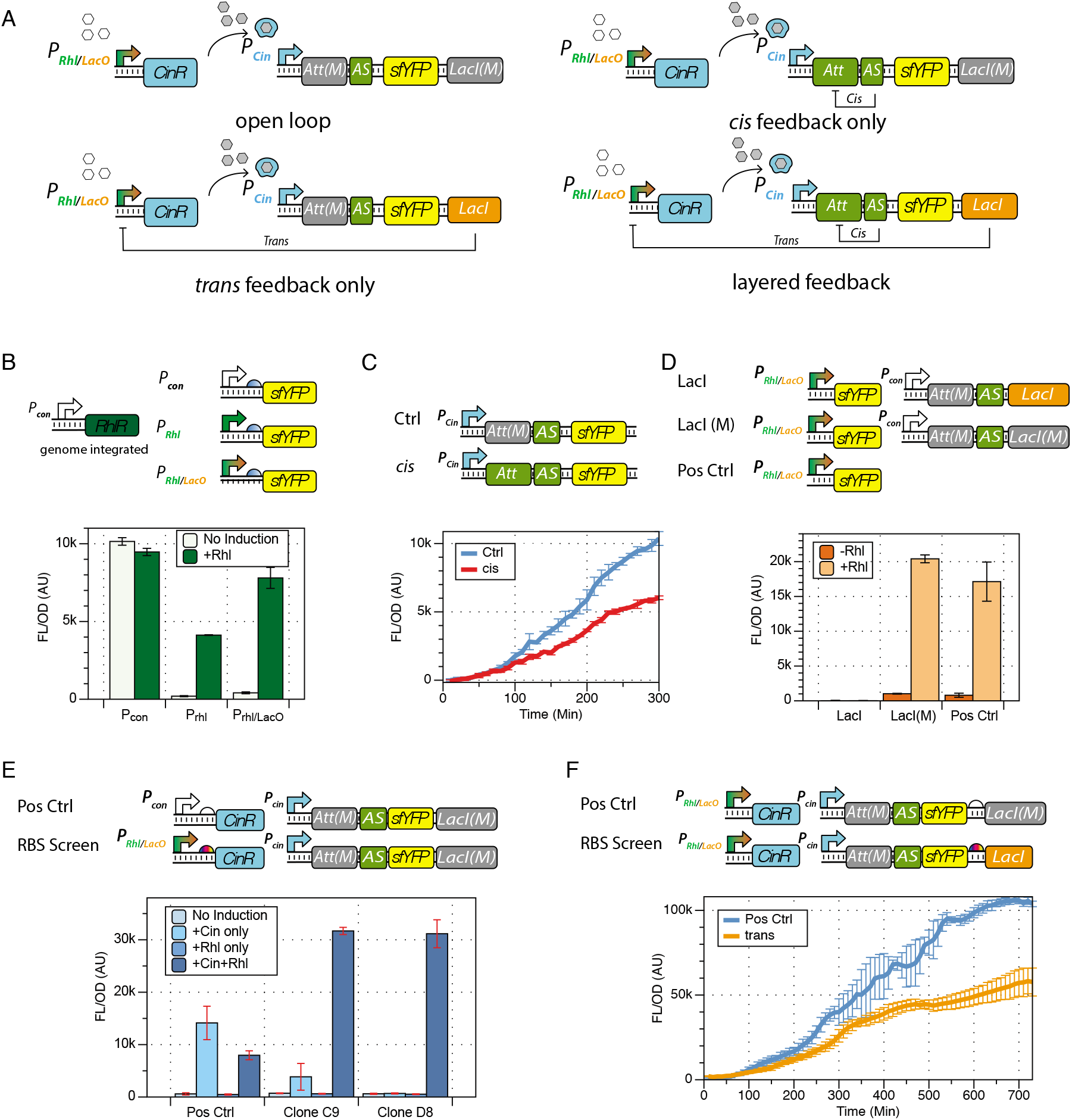
Experimental construction of the layered feedback control in *E. coli*. (A) The four genetic constructs designed for experimental construction of the open loop, the *cis* feedback, the *trans* feedback, and the layered feedback. (B) Partcharacterization of combinatorial promoter *P_Rhl/LacO_*. Induction signal of *P_Rhl/LacO_*, compared with constitutive promoter *P_Rhl_* and Rhl-inducible promoter *P_Rhl_*, performing in RhlR integrated JS006 strain. (C) Part-characterization of sRNA regulator AS with its paired attenuator Att and its orthogonal attenuator Att(M). (D) Part-characterization of combinatorial promoter *P_Rhl/LacO_*, tested with its repressor LacI and a mutated repressor LacI(M). LacI(M) was built off LacI, with the LacO binding site deleted. (E) RBS screening result for the expression of CinR to enable activation cascade. The RBS in Clone D8 was chosen to proceed with the next stages of construction. (F) RBS screening result for the expression of lacI to facilitate the *trans* feedback. The plot shows that the selected clone confirmed the functionality of the negative feedback without over-repression.

To characterize all the parts and confirm their functionalities, we started with the promoter *P_Rhl/LacO_*. Because LacI was designed as a regulatory part of the construct, we chose strain JS006 [26] for its ΔLacI genotype and integrated a constitutively expressed RhlR into its genome. We characterized the promoter *P_Rhl_/_LacO_* in this RhlR containing JS006 strain with 200 *μ*M Rhl induction (Figure 3B). The test confirmed that the combinatorial promoter has sufficient fold induction with about 5% leaky expression. Then, we tested the sRNA mediated *cis* feedback with AS-Att pair and AS-Att(M) pair. We confirmed that the AS-Att(M) pair functions as an appropriate control for the *cis* feedback, as the dynamics of the two constructs are consistent with previous findings [27] (Figure 3C). Next, we examined the repressive regulatory function of combinatorial promoter *P_Rhl/LacO_* with LacI and LacI(M) (Figure 3D). The result confirmed that *P_Rhl/LacO_* can be fully repressed by LacI with full Rhl induction. In the meantime, we determined that LacI(M) is an appropriate control for the *trans* feedback.

At last, we performed two sets of ribosome binding site (RBS) screening assays to achieve the ideal protein expression strength that actuates the desired circuit functionalities. First, we searched for an RBS to express CinR so that it facilitates the activation cascade in our circuit design context. The cascade is designed to be turned on by the inducible expression of CinR through promoter *P_Rhl/LacO_*. Subsequently, CinR binds to inducer Cin in the growth medium, and then the complex activates the promoter *P_cin_*. Because *P_Rhl/LacO_* exhibited some leaky expression, the RBS for CinR required optimization. If the RBS is too weak, the downstream cascade could be under-activated; if the RBS is too strong, the leaky expression of *P_Rhl/LacO_* could cause the cascade to be over-activated without Rhl induction. Here, we screened for an RBS so that the system is off (shows minimal sfYFP signal) with no inducer, only Rhl inducer, or only Cin inducer. Then the system turns on with both Rhl and Cin inducers. As shown in Figure 3E, Clone D8 presented the desired induction profile. The resulting RBS was selected to proceed with the circuit construction. Finally, we screened the RBS for LacI so that it facilitates the *trans* feedback. As we see in Figure 3D, LacI has a strong repressive effect on *P_Rhl/LacO_* with full Rhl induction. Therefore, a relatively weak RBS is required for the *trans* feedback to function. Figure 3F showed the desired dynamical profile of a clone resulting from the screen, where LacI is expressed enough to present a repressed trajectory yet is not over-expressed to shut off the sfYFP expression entirely.

### Interrogating the Dynamics of Layered Feedback Control in *E. coli*

After building the four testing constructs in Figure 3A, we sought to confirm whether our constructs were properly built by comparing measured dynamical profiles with simulated dynamics. Based on the simulated dynamical profile we generated with the reduced model (Figure 4A, Equation 4 to 11), we expected the *trans*, the *cis*, and the layered feedback constructs to have lower equilibrium output signals and faster responses after the induction of Rhl and Cin at *t* = 0. We indeed observed this profile experimentally in the first 7 hours of our dynamical experiment (Figure 4B). However, as the experiment continued to 16 hours, we observed that the reduced model no longer qualitatively describes the observed dynamics (Figure 4C). Notably, the experimental dynamics no longer reach equilibrium. Is this discrepancy due to errors in circuit construction or in dynamical models?

**Figure 4:**
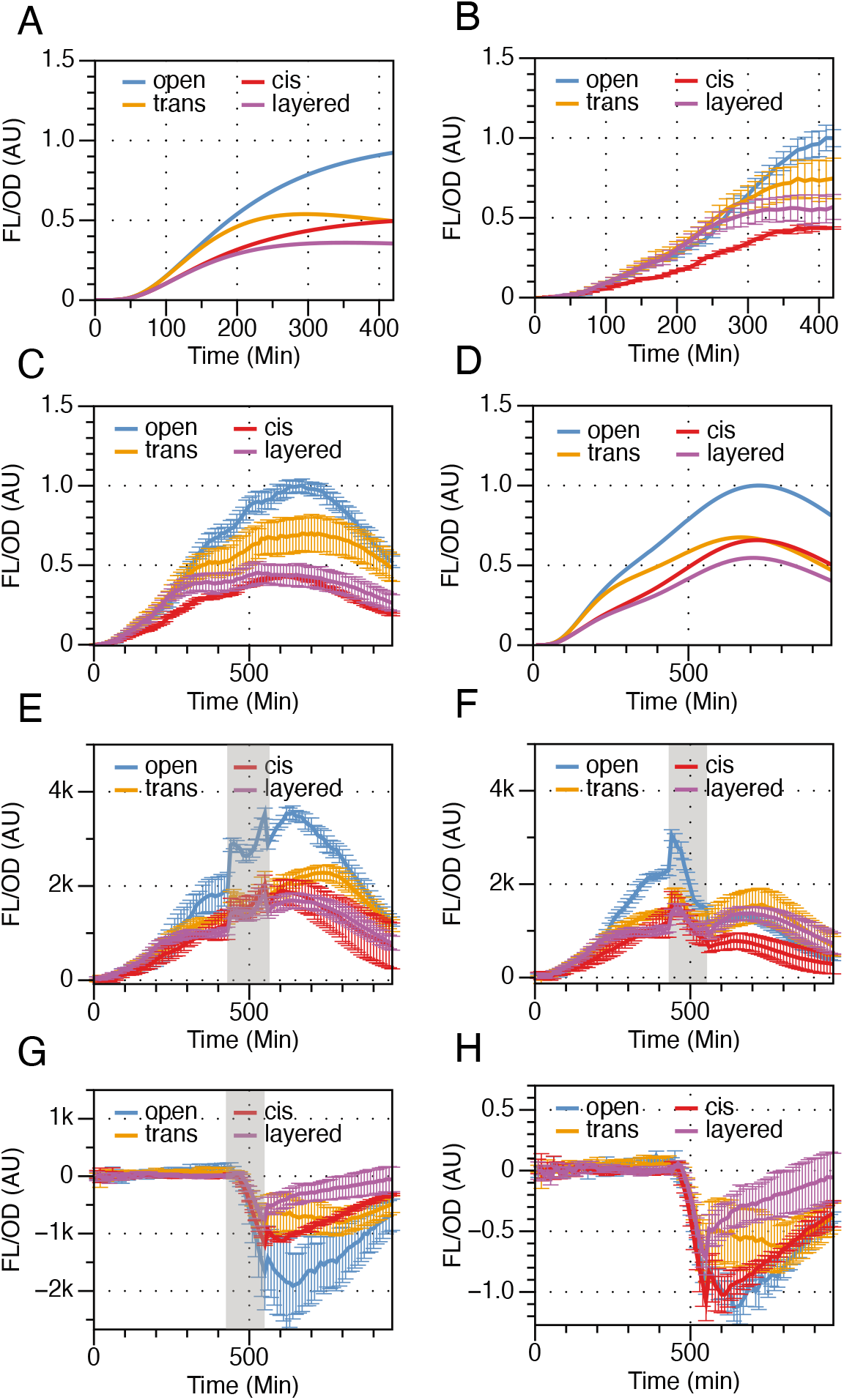
Dynamics of the four testing constructs – the open loop, the *cis* feedback, the *trans* feedback, and the layered feedback, as shown in Figure 3A. (A) Simulated dynamics of the four constructs with reduced model in Equation 4–11 run for 7 hours. (B) Experimentally observed dynamics of the four constructs run for 7 hours. (C) Experimentally observed dynamics of the four constructs run for 16 hours. (D) Simulated dynamics of the four constructs with growth-dependent model in Equation in SI section 1 run for 16 hours. (E)-(H) Dynamic analysis of the test and intact groups in Rhl and Cin wash perturbation experiment. Grey regions indicate the duration of disturbances. (E) Dynamical profile of the intact group. (F) Dynamical profile of the test group. (G) Disturbance profile, calculated by subtracting the dynamics of the test group by the dynamics of the intact group. (H) Scaled disturbance profile. The dynamical deviation of the test group from the intact group, scaled by each construct’s pre-disturbance output value.

We knew that the reduced model was built with numerous assumptions that do not hold as a bacterial culture grows towards capacity. One of many affected parameters is protein dilution *d*. The reduced model assumes a constant cell division rate, therefore a constant d. However, we observed that the culture grew beyond the exponential phase and started to slow down as population reaches capacity (Figure S1). As cell division slows down, protein division slows down, but the active protein degradation due to nutrient starvation speeds up. At the same time, protein production also slow down as it is tied to the nutrient availability in the growth medium. We then expanded our model with functions that approximate the connection between gene expression, cell growth, and nutrient availability (See SI Section 1). The dynamical profile we obtained (Figure 4D) roughly resembles the observation in Figure 4C, which confirmed the functionality of our constructs beyond the exponential phase. Although much more work is needed to create an accurate multiscale model with identified parameters, this preliminary model provided insights for understanding the gene expression dynamics across multiple growth phases.

Since the system dynamics in living cells do not reach equilibrium, we cannot use the same method in Figure 2 to interrogate system dynamics. To overcome this problem, we designed each perturbation experiment with a test group and an intact group. Figure 4E and Figure 4F showed the two groups from one of the eight sets of perturbation experiments. In this experiment, both groups were induced with 100 *μ*L of Rhl and 10 *μ*L of Cin at *t* = 0 and let grow for 7 hours. At hour 7, both groups were washed with phosphate buffered saline (PBS), and the growth medium was replaced. The intact group shown in Figure 4E was resuspended in fresh medium with 100 *μ*L of Rhl and 10 *μ*L of Cin; the test group was resuspended in fresh medium with no inducer. The perturbation window was 120 minutes (greyed section in Figure 4E–H), then both groups were washed and resuspended back to the recycled pre-disturbance media. We measured the disturbance of constructs by the deviation of the test group dynamics from the act group dynamics (Figure 4G). Since the four constructs do not have the same signal output level, we scaled the disturbance profile in Figure 4G by each construct’s pre-disturbance output value to obtain a comparison (Figure 4H). This method allows us to access the dynamical profiles synthetic biological networks under perturbations without a equilibrium. It is also easily applicable to future studies of system amics in biology.

### Evaluating the Robustness-efficiency Performance of Layered Control in *E. coli*

In Figure 5, we show the robustness-efficiency performance of the four constructs in eight different sets of perturbations. These plots were generated by analyzing the scaled disturbance profiles of each culture in perturbation experiments. One of the eight experiments was shown in Figure 4E–4H. Due to the low nutrient availability in the stationary growth phase, systems also do not necessarily adapt perfectly after the perturbation is removed. Therefore, the 40% recovery time (*T*_40_) was used as the efficiency metric, which was defined by the time it took for a system to recover 40% from its peak response after the perturbation was removed (Figure S4A). if a trajectory failed to recover 40%, we took the final time measurement as its *T*_40_. The peak response was measured from the scaled disturbance profiles, one example of which is shown in Figure 4H. We analyzed the replicates individually. Each experimental trajectory was smoothed by an 8 unit moving average before the peak response was identified. For each type of perturbation, we plotted the peak response in the y-axis and the *T*_40_ in the x-axis for all 20 trajectories (5 for each construct). For instance, Figure 5A is the trade-off plot generated with the dynamics shown in Figure 4E-H, with Cin and Rhl wash. The shaded regions marked one standard deviation of the robust-efficiency metrics of each construct. To examine system dynamics with positive perturbation, we also perturbed the system with a Cin and Rhl spike, as shown in figure 5B. Subsequently, we perturbed the system with Rhl only (Figure 5C, 5D), and the growth temperature (Figure 5E, 5F) in both negative and positive directions to evaluate system dynamics under perturbations. In addition, we also perturbed the systems with glucose concentration. However, these trajectories did not trend towards adaptation (Figure S3). Therefore, their efficiency metric was defined by the initial response time of induction (Figure S4B), as shown in Figure 5G, H and Figure S5. The ideal performance region of robust and efficient was highlighted with the yellow color gradient. If a dot on this profile is closer to the highlighted region, it is considered to have a better dynamical performance under a certain perturbation. The detailed protocols of these dynamic experiments are listed in Table S3.

In electrical engineering and control theory, a system’s response to disturbances is usually evaluated with a single perturbed input (Figure 1C). However, in biological systems, perturbations are often applied at the environmental level, which could have much broader, yet poorly understood impacts on the system dynamics. In our node-based design (Figure 1B) and generic biomolecular design (Figure 2A), perturbations were introduced as universal low-frequency transcriptional pulses. Here, we first applied the same perturbation at both negative and positive directions (Figure 5A, 5B) with Cin and Rhl. We saw that although the trade-off profiles of the *trans* and the *cis* feedbacks with the same perturbation in opposite directions were not consistent with each other, the layered feedbacks had advantageous robust-efficiency performance in comparison to the single-layer controls in both cases. It is worth noting that in Figure 5B, the open loop appeared to have the best performance, which was unexpected. We hypothesize that this is caused by endogenous gene expression limitation-induced disturbance attenuation. The open loop constructs have the highest output signal; when the synthetic system was spiked with more inducer to increase the circuit’s transcriptional strength, the endogenous system is at its capacity to up-regulate the system output at the transnational level. Nevertheless, the remaining controlled constructs showed that the layered feedback control inherited performance characteristics from the two single-layer constructs. After that, the next set of disturbances were induced through perturbation of system input Rhl in both directions. In this set of experiments, the *trans* feedback, compared to the *cis* feedback, appears to recover slower after the perturbation. More importantly, the layered control appeared to have a faster response and stronger resilience than the single-layer feedbacks. Finally, in the temperature perturbations that emulate the most common environmental disturbances in biological systems (Figure 5E, F), we perturbed the system by dropping the incubation temperature from 37°C to 30°C (Figure 5E) and spiked it with a temperature increase from 37°C to 42°C (Figure 5F). We observed that systems with layered control performed better in the trade-off plots for both sets of environmental perturbations. In the final set of perturbation experiments, we dropped the glucose concentration in the growth medium from 1% to 0% to apply a negative perturbation and raised it from 0.1% to 1% to apply a positive perturbation (Figure 5G, H). Although they did not trend towards adaptation (Figure S3), the trade-off plots with initial response also showed that systems with layered control were more robust and efficient.

**Figure 5:**
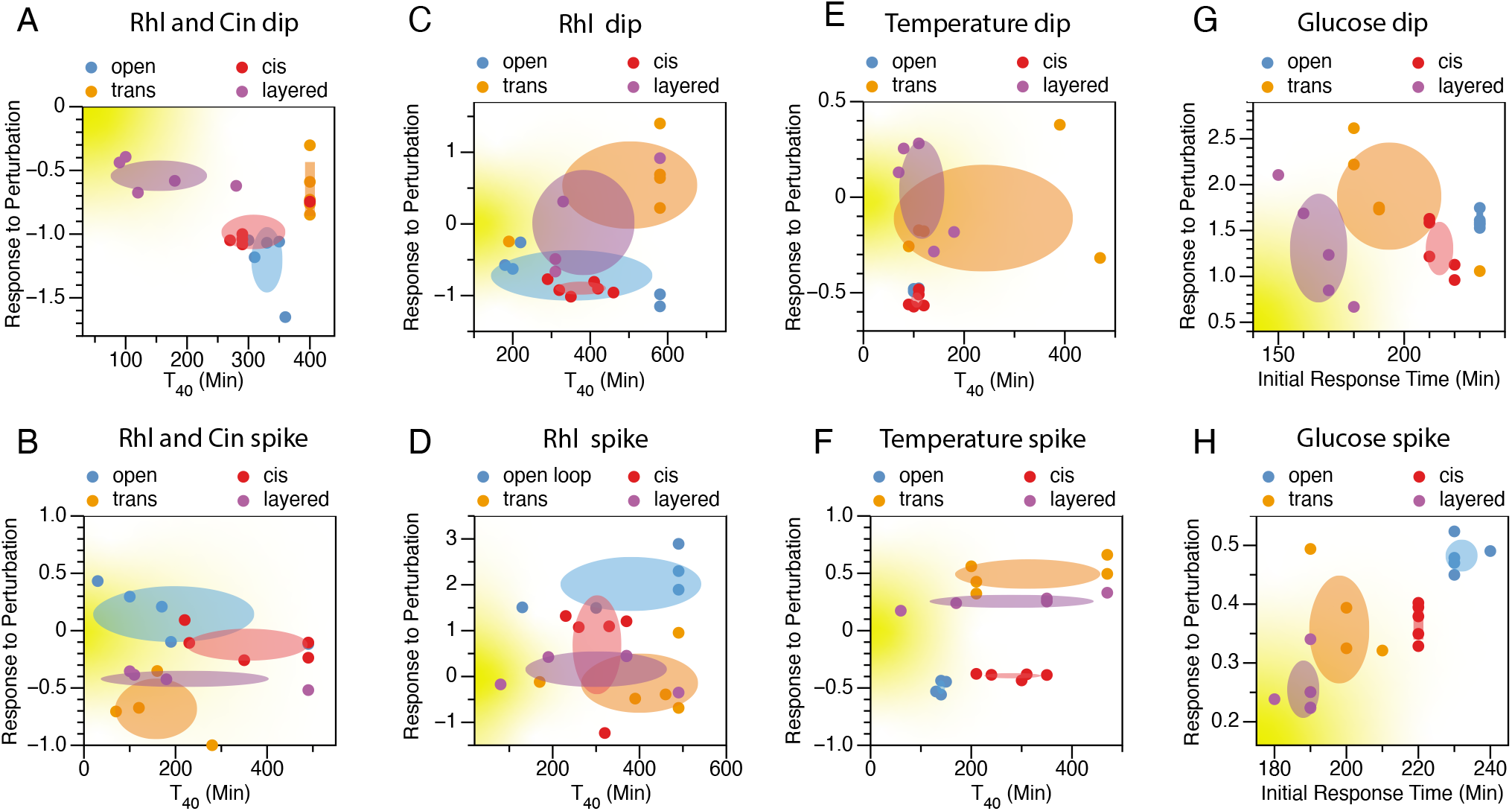
Robustness-efficiency performance of the four constructs in Figure 3A under perturbations presented in trade-off plots. The eight sets of perturbations are (A) Cin and Rhl concentration dip, (B) Cin and Rhl concentration spike, (C) Rhl only concentration dip, (D) Rhl only concentration spike, (E) growth temperature dip from 37°C to 30°C, (F) growth temperature spike from 37°C to 42°C, (G) glucose concentration drop from 1% to 0.1%, (H) glucose concentration spike from 0.1% to 1%. All plots were generated with the scaled disturbance profile (Figure 4H and Figure S2, S3) in each set of the perturbation experiments. For each biological replicate, the peak value in the dynamical profile is plotted on the y-axis to represent its response to perturbations; the time it takes for a trajectory to recover 40% of the peak disturbance is termed *T*_40_, which is plotted in the x-axis in plots (A)-(F) (Analysis methods shown in Figure S4A). Because the dynamical profiles of glucose disturbances did not trend towards adaptation (shown in Figure S3), the initial response time was plotted on the x-axis in (G) and (H) as the efficiency metric. In all plots, each dot renders the robustness-efficiency performance of individual cell culture; each shaded area represents the robustness-efficiency performance of a construct within one standard deviation. The yellow gradient shaded area indicate the ideal performance (robust and efficient). An ideal dynamical system is robust (with a low response to perturbation) and efficient (with a small *T*_40_ value or initial response time).

Through the measurement and analysis of all eight sets of experiments, many aspects of the dynamics posed interesting questions regarding the natural biological networks. For instance, when given a glucose wash, all four constructs exhibited an increase in gene expression, and this increase was sustained long after the disturbance was removed (Figure S3). Moreover, when the incubating temperature was raised from 37°C to 42°C, the *trans* and the layered feedback constructs demonstrated positive disturbances while *cis* feedback and open loop demonstrated negative disturbances (Figure S3). Interestingly, this observation was reversed when the systems were perturbed with Rhl and Cin spikes. The *trans* feedback and the layered feedback showed negative disturbances, while the *cis* feedback and the open loop demonstrated positive disturbances (Figure S2). In the Rhl dip perturbation experiment, although all four constructs revealed an output signal dip during the perturbation window, their dynamics in the recovery window progressed in different directions (Figure S2). These observations suggested that pieces in our synthetic circuits were coupled through unknown mechanisms that await discoveries, which provided new directions for future works. Nevertheless, without knowing all the mechanistic details, we were able to experimentally confirm that layered feedback overcomes the robustness-efficiency trade-off limit that constrains single-layer controls and achieves better control performance.

## Conclusion

This work provided experimental proof that layered feedback control as a control optimization strategy is applicable in biomolecular networks. We first showed that the layered architecture overcomes the robustness-efficiency trade-off in a wide range of parameter space with a node-based minimal model. Then we expanded the design to a generic biomolecular model to simulate system dynamics with perturbations. With the guidance of theory and simulation, we forward engineered a layered feedback controlled synthetic network in living *E. coli* cells. We successfully constructed this network with a combinatorial regulation, a activating cascade, and two nested autoregulatory motifs to achieve its desired dynamical functions. Finally, in eight sets of different dynamical perturbation experiments, we observed that the layered control feedback consistently overcomes the robustness-efficiency trade-off limit. This result not only provides validated theory guidance for controller design for synthetic biological networks. It also offers insights for understanding nature’s dynamical control strategies.

The experimentally observed gene expression dynamics across different growth phases also revealed the dynamical entanglement across the populational and the molecular scales. We found that the common assumptions on constant cell division and unlimited resources are responsible for the major discrepancies between simulated and observed system dynamics. For system modeling, the quality of a model is bound by the accuracy and identifiability trade-off. Reduced models have fewer species and parameters so that they are more identifiable but less accurate and relevant due to over-simplification. On the other hand, chemical reaction network models are more accurate with fewer assumptions, but they are usually substantial in size and difficult to system identify. For future studies, we plan to layer the two types of models to overcome this trade-off. Also, we will expand the utilization of experimental data for system identification. We hope to utilize current collected dynamical data from this work to improve the accuracy and utility of biomolecular models and discover unknown mechanistic details in biomolecular networks.

## Methods and Materials

### Plasmid Construction and Purification

All plasmids used in this study were created using Golden Gate [28] assembly, Gibson assembly, or 3G assembly [29], with NEB^®^ Turbo Competent *E. coli* as cloning strain. Plasmids were purified using a Qiagen QIAprep Spin Miniprep Kit (Qiagen 27104). Linear fragments were gel extracted and purified using MinElute Gel Extraction Kit (Qiagen 28606).

### Model Simulations

All models were simulated by solving corresponding ODEs using MATLAB function ode15s over a set of discrete time steps using guessed or estimated parameters.

### Strains, Growth Media and In-cell Part Characterization Experiments

All experiments were performed in *E. coli* strain JS006 [26] with either constitutively expressed RhlR or constitutively expressed CinR and RhlR [23] integrated into the chromosome with pOSIP integration plasmid into O-site containing Kanamycin resistance gene [30]. Plasmids with different constructs were transformed into the modified JS006 competent cells. Depending on the antibiotic resistance plated on LB + Agar plates containing 100 *μ*g/ml carbenicillin or/and chloramphenicol, and incubated overnight at 37°C. At least three colonies of each experimental condition were inoculated into 200 *μ*L of LB containing carbenicillin in a 2 mL 96-well block, and grown overnight at 37°C at 1000 rpm in a bench top shaker. Four microliters of the overnight culture were added to 196 *μ*L of M9 supplemented M9 media (1X M9 minimal salts, 0.5 *μ*g/mL thiamine hydrochloride, 1.0% glucose, 0.1% casamino acids, 0.2 mM *MgSO_4_*, 0.1 mM *CaCl_2_*) containing carbenicillin and/or chloramphenicol and grown for 4 hours at 37°C at 1000 rpm.

For single time-point measurements (Figure 3B, D, E), the sub-culture was then diluted 20 times with M9 its appropriate antibiotics and AHL inducers and grown in a 96 well plate for 6 hours before the measurement of fluorescence (504 nm excitation, 540 nm emission) and optical density (OD, 600 nm). The two AHL inducers used in this work are N-(3-Oxotetradecanoyl)-L-homoserine lactone (Cin, Sigma O9264) and 3-Hydroxy-C4-HSL, N-(3-Hydroxybutanoyl)-L-homoserine lactone (Rhl, Sigma 74359). Experiment in Figure 3B was grown with chloramphenicol, containing 0 *μ*M or 200 *μ*M of Rhl inducer; Experiment in Figure D was grown with carbenicillin and chloramphenicol, containing 0 *μ*M or 200 *μ*M of Rhl inducer; Experiment in Figure 3E was grown with carbenicillin and chloramphenicol with four inducer conditions: 0 *μ*M Cin and 0 *μ*M Rhl, 10 *μ*M Cin and 0 *μ*M Rhl, 0 *μ*M Cin and 200 *μ*M Rhl, and 10 *μ*M Cin and 200 *μ*M Rhl.

For time-course characterization measurements (Figure 3C, F), the sub-culture was then diluted 20X to 200 *μ*L with M9 its appropriate antibiotics and inducers and on a 96 well plate in a Biotek SynergyH1 plate reader. The experiment in Figure 3C was grown in M9 media with carbenicillin and 10 *μ*M Cin for 5 hours. Experiment in Figure 3F was grown in M9 media with carbenicillin, 10 *μ*M Cin and 200 *μ*M Rhl for 12 hours. The plate reader incubates the runs at 37°C with maximum linear shaking. It measures fluorescence (504 nm excitation, 540 nm emission) and optical density (OD 600 nm) every 10 minutes.

For RBS screening experiments that resulted in the appropriate constructs shown in figure 3D and 3E, the plasmids were assembled with an ARL (Anderson RBS library) using 3G assembly [29]. Colonies are picked and grown in 200 *μ*L LB with carbenicillin and chloramphenicol overnight at 37°C at 1000 rpm on a benchtop shaker. Four microliters of the overnight culture were added to 196 *μ*L of M9 supplemented M9 media to grow to the exponential phase before the experiment. For the RBS screen experiment on CinR expression (Figure 3E), each culture resulted from a single colony was diluted 20X and grown in M9 media, induced with either Rhl only, or Cin and Rhl. The colonies that appeared to be “off” with Rhl and “on” with Cin and Rhl were selected. For the RBS screen experiment on LacI expression (Figure 3F), each culture resulted from a single colony was diluted 20X and grown in M9 media, induced with Rhl and Cin. Along with the screened colonies, a triplicate of positive control with LacI(M) was measured to provide screening reference (Figure 3F). The colonies with signals that are apparent but lower than the positive control references were selected. All selected colonies for RBS screening were sequenced, the resulting RBS sequenced were cloned into the constructs for verification with at least three replicates.

### Dynamical Perturbation Experiments

All perturbation experiments were performed in JS006 *E.coli* with genome integrated RhlR. A blank ColE1 high copy plasmid is transformed with the *P_rhl/lacO_* controlled cassette on p15A backbone to serve as the negative control. Four colonies of negative control and five colonies of each of the four testing constructs showed in Figure 3A were picked. Cultures were grown in 200 *μ*L LB with carbenicillin and chloramphenicol overnight at 37°C at 1000 rpm on a benchtop shaker. Four microliters of the overnight culture were added to 196 *μ*L of M9 media and grown for four fours to prepare the sub-culture. The experiments started with a 20X dilution of the sub-culture into M9 media with carbenicillin and chloramphenicol, induced with Cin and Rhl. The plate reader incubates the runs at 37°C with maximum linear shaking. It measures fluorescence (504 nm excitation, 540 nm emission) and optical density (OD, 600 nm) every 10 minutes. The first stage of the run lasts 5.5-7 hours. After disturbances were introduced, both intact and perturbed cultures were grown in the plate reader for 2-3 hours, and the fluorescent and OD dynamics were collected. Then, the disturbance of both groups were removed. Subsequently, the run continued in the plate reader for the remainder of the 16 hours. The detailed protocols of each perturbation are listed in Table S3.

## Acknowledgement

The authors would like to thank John Doyle, Fangzhou Xiao, Ayush Pandey, and Xinying Ren for their insightful discussions. The author C. Y. Hu is partially supported by Defense Advanced Research Projects Agency (Agreement HR0011-17-2-0008). The content of the information does not necessarily reflect the position or the policy of the government, and no official endorsement should be inferred.

