## Supplemental Info for "Layered Feedback Control Overcomes Performance Trade-off in Synthetic Biomolecular Networks"

Chelsea Y. Hu[1] and Richard M. Murray[1]

[1]Division of Biology and Biological Engineering, California Institute of Technology, Pasadena, CA.

### 1 Model of the coupled dynamics of gene expression and cell growth

To model the biomolecular dynamics coupled with cell growth, we wrote the following equations of 15 species and 25 parameters. Species 1-14 are in the scale of single cell. The last species C is in the populational.

#### 1.1 Equations

$$\frac{dM_{cin}}{dt} = \boldsymbol{\beta_{rhl}} \cdot (\frac{X_{rhl}}{X_{rhl} + K_{rhl}} + l_{rhl}) \cdot f_{trans} - \boldsymbol{d_m} \cdot M_{cin} + k_r \cdot T_{cin} - M_{cin} \cdot B \cdot k_{tf} \tag{1.1}$$

$$\frac{dP_{cin}}{dt} = k_r \cdot T_{cin} - m_r \cdot P_{cin} - \boldsymbol{d_g} \cdot P_{cin} - \boldsymbol{d_p} \cdot P_{cin} \tag{1.2}$$

$$\frac{dC_{ind}}{dt} = m_r \cdot P_{cin} - \boldsymbol{d_g} \cdot C_{ind} - \boldsymbol{d_p} \cdot C_{ind} \tag{1.3}$$

$$\frac{dR}{dt} = \boldsymbol{\beta_{cin}} \cdot (\frac{C_{ind}}{C_{ind} + K_{cin}} + l_{cin}) \cdot f_{cis} - \boldsymbol{d_r} \cdot R - m_{as} \cdot R \tag{1.4}$$

$$\begin{aligned} \frac{dM}{dt} &= \boldsymbol{\beta_{cin}} \cdot (\frac{C_{ind}}{C_{ind} + K_{cin}} + l_{cin}) \cdot f_{cis} - \boldsymbol{d_m} \cdot M + k_r \cdot T_{lac} - M \cdot B \cdot k_{tf} + T_{lac} \cdot k_{lac_R} \\ &+ k_r \cdot T_{fp} - M \cdot B \cdot k_{tf} + T_{fp} \cdot k_{fp_R} \end{aligned} \tag{1.5}$$

$$\frac{dP_{lac}}{dt} = k_r \cdot T_{lac} - \boldsymbol{d_g} \cdot P_{lac} - \boldsymbol{d_p} \cdot P_{lac} - m_{lac} \cdot P_{lac} \tag{1.6}$$

$$\begin{aligned} \frac{dB}{dt} &= -M_{cin} \cdot B \cdot k_{tf} + T_{cin} \cdot k_{cin_R} - 2 \cdot M \cdot B \cdot k_{tf} + T_{lac} \cdot k_{lac_R} + T_{fp} \cdot k_{fp_R} \\ &+ k_r \cdot (T_{cin} + T_{lac} + T_{fp}) - \boldsymbol{d_p} \cdot B \end{aligned} \tag{1.7}$$

$$\frac{dP_{fp}}{dt} = k_r \cdot T_{fp} - \alpha \cdot P_{fp} - \boldsymbol{d_g} \cdot P_{fp} - \boldsymbol{d_p} \cdot P_{fp} \tag{1.8}$$

$$\frac{dT_{cin}}{dt} = -k_r \cdot T_{cin} + M_{cin} \cdot B \cdot k_{tf} - T_{cin} \cdot k_{cin_R} - \boldsymbol{d_g} \cdot T_{cin} \tag{1.9}$$

$$\frac{dT_{lac}}{dt} = -k_r \cdot T_{lac} + M \cdot B \cdot k_{tf} - T_{lac} \cdot k_{lac_R} - \boldsymbol{d_g} \cdot T_{lac} \tag{1.10}$$

$$\frac{dT_{fp}}{dt} = -k_r \cdot T_{fp} + M \cdot B \cdot k_{tf} - T_{fp} \cdot k_{fp_R} - \boldsymbol{d_g} \cdot T_{fp} \tag{1.11}$$

$$\frac{dR_m}{dt} = m_{as} \cdot R - \boldsymbol{d_r} \cdot R_m \tag{1.12}$$

$$\frac{dP_{m_{lac}}}{dt} = m_{lac} \cdot P_{lac} - \boldsymbol{d_g} \cdot P_{m_{lac}} - \boldsymbol{d_p} \cdot P_{m_{lac}} \tag{1.13}$$

$$\frac{dP_{fp}}{dt} = \alpha \cdot P_{fp} - \boldsymbol{d_g} \cdot P_{m_{fp}} - \boldsymbol{d_p} \cdot P_{fp} \tag{1.14}$$

$$\frac{dC}{dt} = r_g \cdot (1 - \frac{C}{C_{max}}) \cdot C \tag{1.15}$$

Note that the termed in red ink describes the translational mechanism of the three protein species: $CinR$, $lacI$, and sfYFP. The chemical reactions are described by the following chemical equation:

$$Ribosome(B) + mRNA(M_x) \underset{k_{xR}}{\overset{k_{tf}}{\rightleftharpoons}} Tanscript(T_x) \xrightarrow{k_r} Protein(P_x)$$

Also, to connect gene expression with resource limitation, we wrote the bolded parameters as functions of growth and total species in a cell. Specifically, proteins are diluted ($d_g$) by cell division, which slows down when population reaches density capacity. At the same time, the degradation rates of proteins ($d_p$) are also functions of growth, which speeds up when cell population reaches density capacity as the resources are limited and dead cells become the main resource of amino acids in the growth medium; and slow down when there is abundance of total proteins($P_{tot}$). The transcriptional rates $\beta_{rhl}$ and $\beta_{cin}$ are also modeled as functions of cell population and species population. Finally, we modeled the degradation rate of RNA species as a function of total RNA in the cell (R$_{tot}$).

$$
\begin{aligned}
d_g &= r_g \cdot (1 - \frac{C}{C_{max}}) && (1.16)\\
d_p &= d_{p_{max}} \cdot \frac{C}{C_{max}} \cdot (1 - \frac{P_{tot}}{P_{max}}) && (1.17)\\
\beta_{rhl} &= \beta_{rhl_{max}} \cdot (1 - \frac{R_{tot}}{R_{max}}) \cdot (1 - \frac{C}{C_{max}}) && (1.18)\\
\beta_{cin} &= \beta_{cin_{max}} \cdot (1 - \frac{R_{tot}}{R_{max}}) \cdot (1 - \frac{C}{C_{max}}) && (1.19)\\
d_r &= d_{r_{max}} \cdot (1 - \frac{R_{tot}}{R_{max}}) && (1.20)\\
d_m &= d_{m_{max}} \cdot (1 - \frac{R_{tot}}{R_{max}}) && (1.21)\\
R_{tot} &= M_{cin} + R + M + T_{cin} + T_{lac} + T_{fp} + R_m && (1.22)\\
P_{tot} &= P_{cin} + C_{ind} + P_{lac} + B + P_{fp} + P_{m_{lac}} + P_{fp} && (1.23)
\end{aligned}
$$

Additionally, $f_{cis}$ and $f_{trans}$ denote for the *cis* and the *trans* feedbacks in the system.
For the open loop: $f_{cis} = 1; f_{trans} = 1$
For the *cis* feedback only: $f_{cis} = \frac{K_R}{K_R+R_m}; f_{trans} = 1$
For the *trans* feedback only: $f_{cis} = 1; f_{trans} = \frac{K_{lac}}{K_{lac}+P_{m_{lac}}}$
For the layered feedback: $f_{cis} = \frac{K_R}{K_R+R_m}; f_{trans} = \frac{K_{lac}}{K_{lac}+P_{m_{lac}}}$

#### 1.2 Model species and parameters

Table S1: Growth Dependent Biomolecular Model Species

| Species | Description |
|---|---|
| $M_{ind}$ | mRNA of CinR |
| $P_{ind}$ | CinR translated peptides |
| $C_{ind}$ | CinR activating complex, with folded CinR bonded with Cin-AHL molecules |
| $R$ | antisense sRNA repressor unfolded |
| $M$ | mRNA containing sRNA, sfYFP and LacI |
| $P_{lac}$ | LacI protein translated peptides |
| $P_{fp}$ | sfYFP protein translated peptides |
| $B$ | Ribosome |
| $T_{cin}$ | Ribosome bound CinR mRNA |
| $T_{lac}$ | Ribosome bound LacI mRNA |
| $T_{fp}$ | Ribosome bound sfYFP mRNA |
| $R_m$ | folded mature antisense sRNA |
| $P_{m_{lac}}$ | folded mature LacI protein |
| $P_{m_{fp}}$ | folded mature sfYFP |
| $C$ | Number of cells in the culture |

Table S2: Growth Dependent Biomolecular Model Parameters

| Parameters | Description | Unit | Estimates |
|---|---|---|---|
| $\beta_{rhl_{max}}$ | Max transcription rate of inducible promoter $P_{rhl/LacO}$ | nM/min | 5 |
| $K_{rhl}$ | Activation coefficient of inducer Rhl | nM | 1.4e5 |
| $K_R$ | Repression coefficient of the sRNA repressor | nM | 20 |
| $d_{m_{max}}$ | Max degradation/dilution rate of mRNA | 1/min | 0.2 |
| $k_{cin_R}$ | Unbinding rate of ribosome B and cinR mRNA | 1/min | 3000 |
| $l_{rhl}$ | Leak coefficient of $P_{rhl/LacO}$ promoter | N/A | 0.10 |
| $m_r$ | Maturation rate of CinR-AHL complex | 1/min | 0.05 |
| $K_{cin}$ | Repression coefficient of sRNA repressor | nM | 80 |
| $\beta_{cin_{max}}$ | Max transcription rate of $P_{cin}$ promoter inducible by $C_{ind}$ | nM/min | 20 |
| $d_{r_{max}}$ | Max degradation/dilution rate of sRNA | 1/min | 0.15 |
| $m_{fp}$ | Maturation rate of sfYFP | 1/min | 0.01 |
| $x_{rhl}$ | The activating inducer for $P_{rhl/LacO}$ | nM | 3e5 |
| $k_{fp_R}$ | Unbinding rate of ribosome B and sfYFP mRNA | 1/min | 1000 |
| $k_{tf}$ | Forward binding rate of mRNA and Ribosome | 1/min | 1 |
| $K_r$ | Translate and release rate of activated transcript | 1/min | 0.015 |
| $k_{lac_R}$ | Unbinding rate of ribosome B and LacI mRNA | 1/min | 3000 |
| $K_{lac}$ | Repression coefficient of LacI | nM | 30 |
| $d_{p_{max}}$ | Max protein degradation rate | 1/min | 0.007 |
| $r_g$ | Max growth rate of cells | 1/min | 0.012 |
| $C_{max}$ | Population capacity | 1/min | 8e8 |
| $m_{AS}$ | Maturation rate of sRNA | 1/min | 0.02 |
| $m_{lac}$ | Maturation rate of LacI protein | 1/min | 0.05 |
| $R_{max}$ | RNA capacity in a cell | 1/min | 10000 |
| $P_{max}$ | Protein capacity in a cell | 1/min | 10000 |
| $l_{cin}$ | Leak coefficient of $\mathrm{P}_{cin}$ | N/A | 0.1 |

### 2 Dynamical perturbation experiment

#### 2.1 Supplemented experimental results

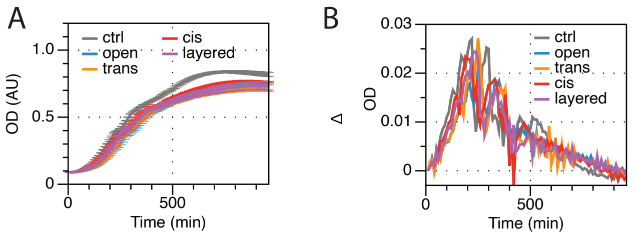

Figure S1: Growth rate is not constant in circuit dynamic window. (A) The OD dynamics for experiment shown in Figure 4C (B) The rate of growth over time for experiment in Figure 4C.

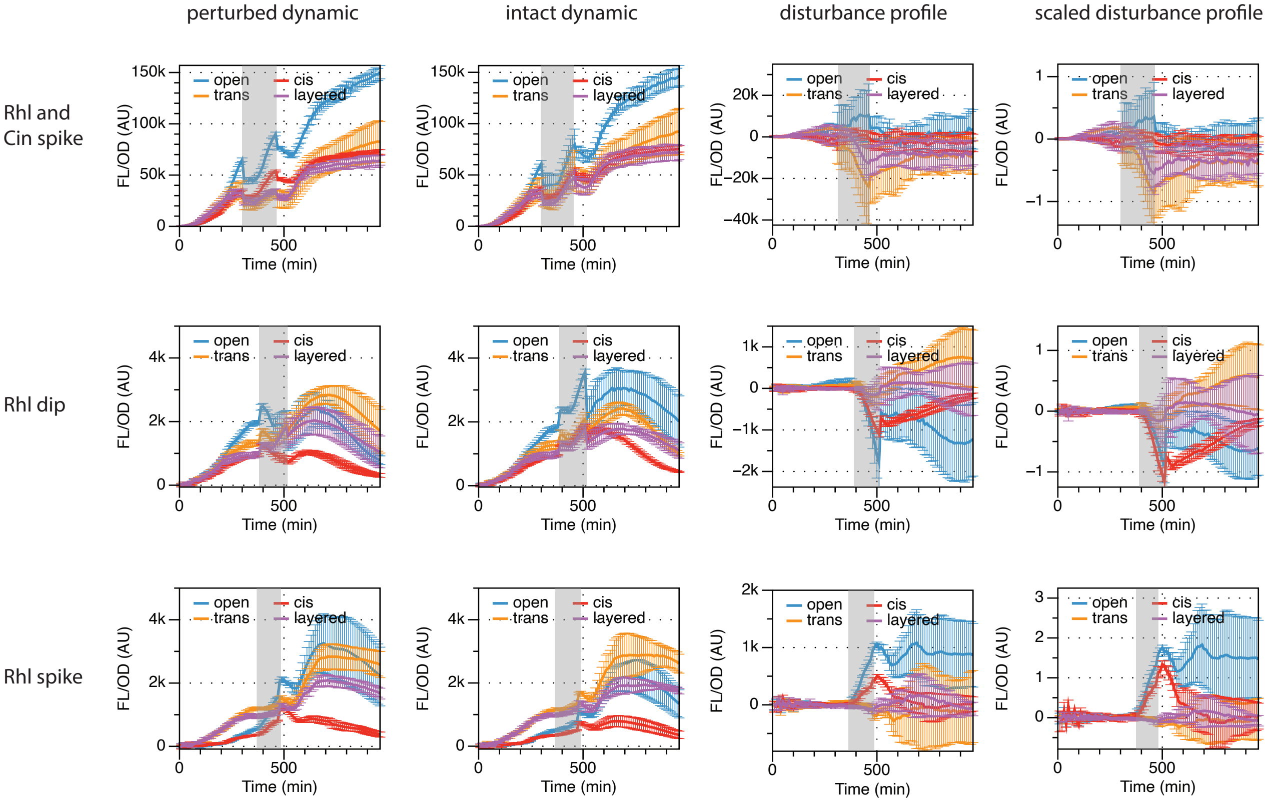

Figure S2: Results of dynamical perturbation experiments, grey regions indicate the duration of perturbations. The three rows correspond to the three different types of perturbations showed in Figure 5B, 5C, and 5D, as listed on the left. The four columns correspond to the dynamics of the perturbed groups, intact groups, disturbance profile (the dynamical deviation of the perturbed group from the intact group), and the scaled disturbance profile (calculated with the disturbance profile scaled by each construct's pre-disturbance output values).

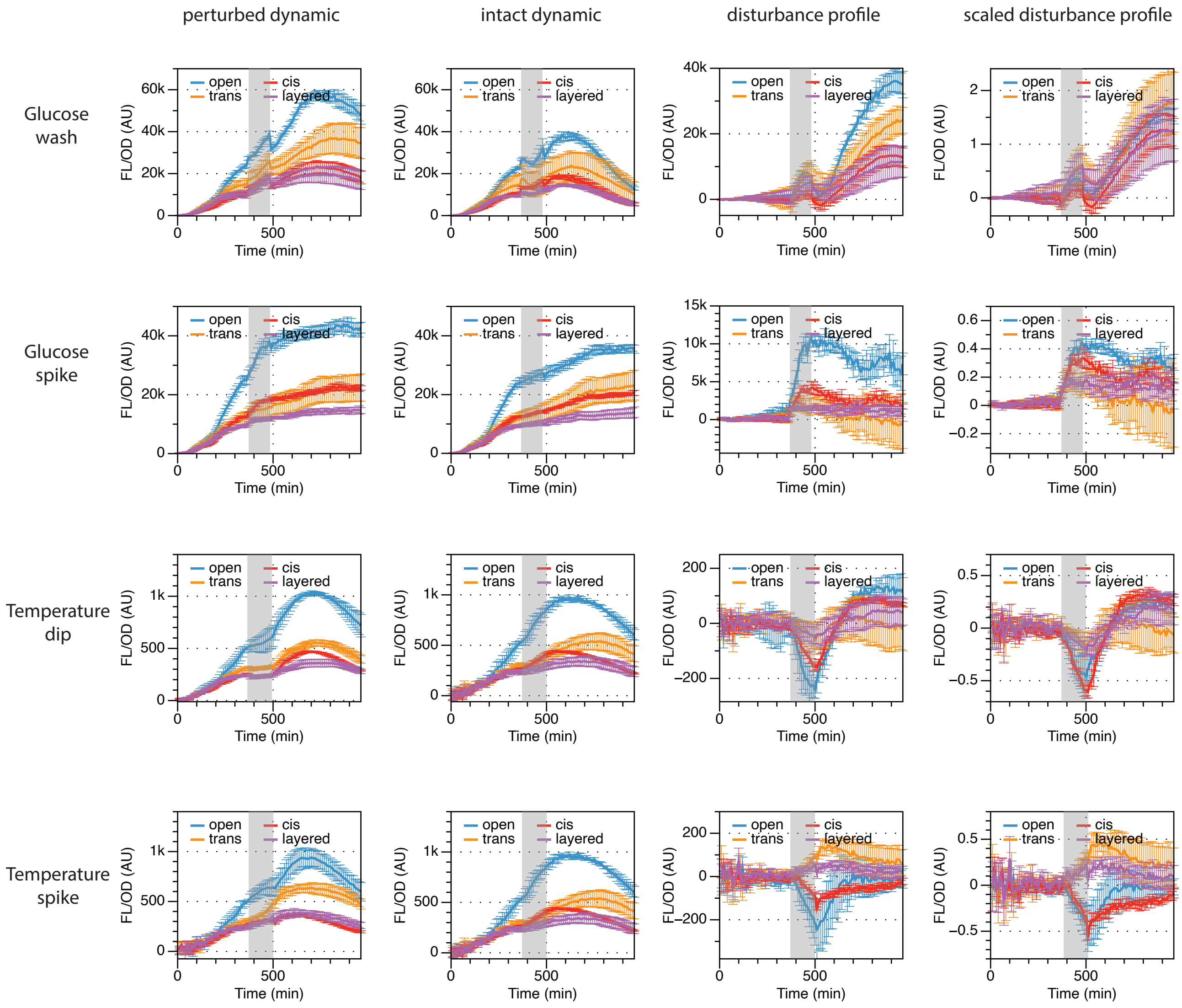

Figure S3: Results of dynamical perturbation experiments, grey regions indicate the duration of disturbances. The four rows correspond to the four different types of perturbations showed in Figure 5E, 5F, 5G, and 5H, as listed on the left. The four columns correspond to the dynamics of the perturbed groups, intact groups, disturbance profile (the dynamical deviation of the perturbed group from the intact group), and the scaled disturbance profile (calculated with the disturbance profile scaled by each construct's pre-disturbance output values).

#### 2.2 Robustness and efficiency analysis on experimental measurements

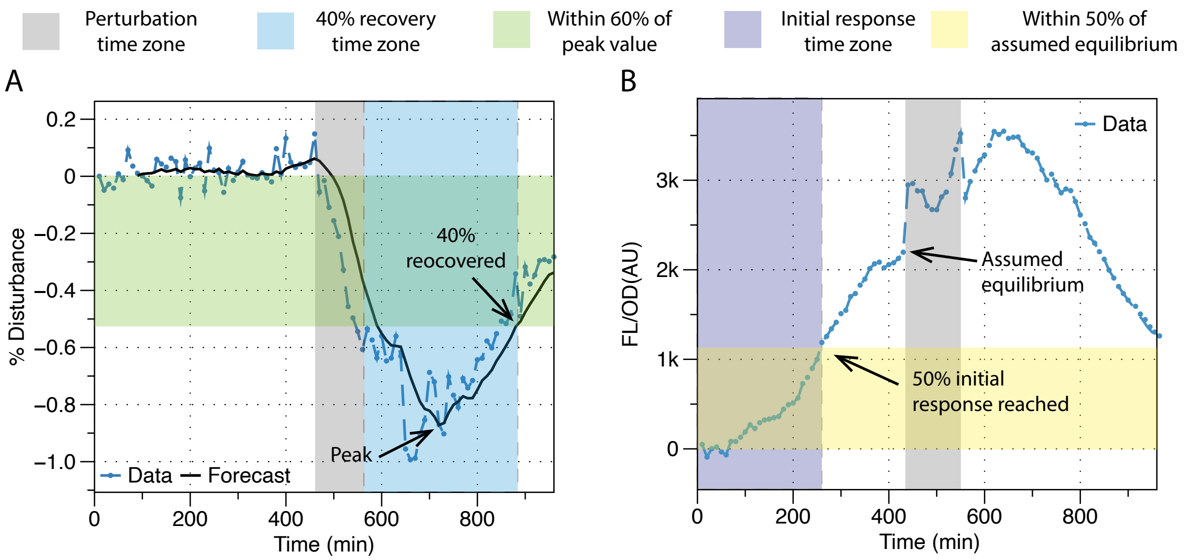

Figure S4: Robustness and efficiency analysis on experimental measurements. (A) Metrics used to generate Figure 5(A)-(F). The blue dash line is the scaled disturbance profile of a single colony taken from Figure 4H. The solid line is a moving average forecast of the experimental data in order to generate a smooth line for analysis. The grey time zone indicates the duration of the perturbation; the blue time zone is the 40% recovery time zone ($T_{40}$), which is calculated from the end of the perturbation to the time it takes to recover 40% of the peak disturbance. The peak disturbance is marked in the plot. The green shaded area is within 60% of the peak value. $T_{40}$ is marked where the green and blue shaded areas intersect. (B) Efficiency metrics used to generate Figure 5(G)-(H) and Figure S5. The blue line is the measured experimental data taken from one of the open loop colonies in Figure 4. The grey time zone indicates the duration of the perturbation; the purple time zone is the initial response time, which is calculated from the system induction time to when the trajectory reaches 50% of the assumed equilibrium. Because systems do not reach equilibrium, this assumed equilibrium is approximated by the last measurement before the perturbation.

#### 2.3 Trade-off plots with initial response time as the efficiency metric

#### 2.4 Detailed experiment protocols

The eight sets of perturbation experiments were performed with the protocol listed in Table S3. For each set of experiment, an intact (Ctrl) group and a perturbed (Test) group were used. In order to isolate the impact of a desired perturbation, both groups undergo the same perturbations that are associated with experimental procedures, i.e. PBS wash, temperature change associated with perturbation introduction, or replenishment of nutrients due to the use of fresh media.

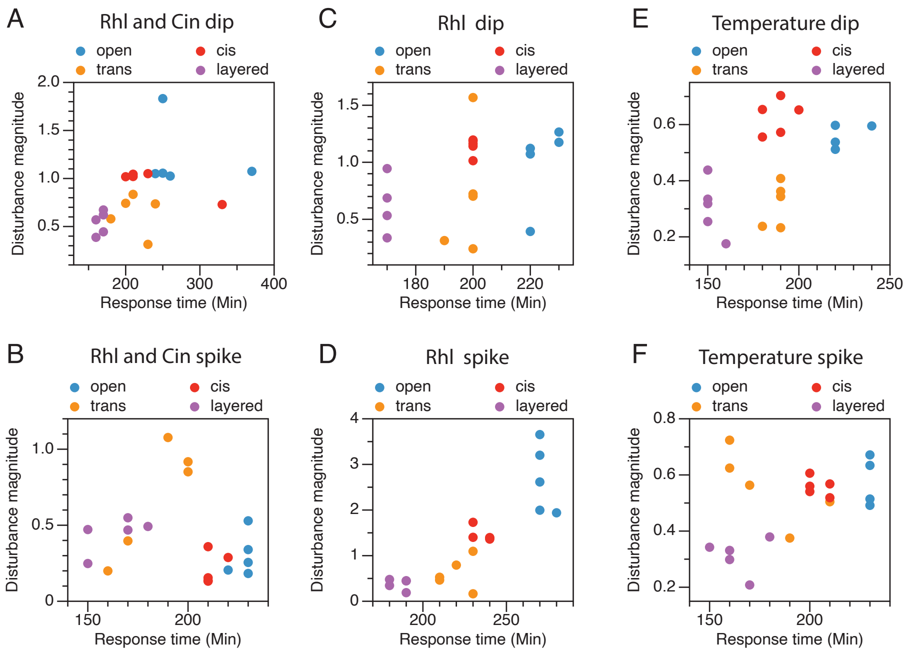

Figure S5: Trade-off plots generated with the initial response time as the efficiency metric. The six sets of perturbations are: (A) Cin and Rhl concentration dip, (B) Cin and Rhl concentration spike, (C) Rhl only concentration dip, (D) Rhl only concentration spike, (E) growth temperature dip from 37°C to 30°C, (F) growth temperature spike from 37°C to 42°C. These plots are generated with the same scaled disturbance profiles that were used to generate Figure 5 (A)-(E).

Table S3: Perturbation experiment protocols

| Experiment | Int. cond. | disturb at | disturbance condition | recover at | recovery cond. |
|---|---|---|---|---|---|
| Rhl+Cin dip Test | standard* | 440 min | wash, standard -Rhl -Cin | 560 min | pre-disturbance media |
| Rhl+Cin dip Ctrl | standard | 440 min | wash, standard fresh | 560 min | pre-disturbance media |
| Rhl+Cin spike Test | low ind.** | 310 min | wash, standard fresh | 470 min | wash, fresh low ind. |
| Rhl+Cin spike Ctrl | low ind. | 310 min | wash, standard fresh | 470 min | wash, fresh low ind. |
| Rhl dip Test | standard | 390 min | wash, standard -Rhl | 520 min | pre-disturbance media |
| Rhl dip CtrlL | standard | 390 min | wash, standard | 520 min | pre-disturbance media |
| Rhl spike Test | standard -Rhl | 360 min | continuous, +Rhl | 480 min | pre-disturbance media |
| Rhl spike Ctrl | standard -Rhl | 360 min | continuous | 480 min | pre-disturbance media |
| Glucose dip Test | standard | 370 min | wash, standard -glucose | 490 min | pre-disturbance media |
| Glucose dip Ctrl | standard | 370 min | wash, standard | 490 min | pre-disturbance media |
| Glucose spike Test | 0.1% glucose | 370 min | continuous, 1% glucose | 490 min | pre-disturbance media |
| Glucose spike Ctrl | 0.1% glucose | 370 min | continuous | 490 min | pre-disturbance media |
| Temp dip Test | standard | 360min | continuous at 30°C | 500 min | continuous standard |
| Temp dip Ctrl | standard | 360min | continuous | 500 min | continuous standard |
| Temp spike Test | standard | 360min | continuous at 42°C | 500 min | continuous standard |
| Temp spike Ctrl | standard | 360min | continuous | 500 min | continuous standard |

** $20\mu M$ Rhl and $3\mu M$ Cin

* M9 medium with 1% glucose, induced with $100\mu M$ Rhl and $10\mu M$ Cin growing at 37°C
